# Capacitation promotes a shift in the energy metabolism in murine sperm

**DOI:** 10.1101/2022.05.08.490683

**Authors:** Maximiliano Tourmente, Ester Sansegundo, Eduardo Rial, Eduardo R. S. Roldan

**Affiliations:** Department of Biodiversity and Evolutionary Biology, Museo Nacional de Ciencias Naturales (CSIC). 28006 Madrid. Spain; Centro de Biología Celular y Molecular, Facultad de Ciencias Exactas, Físicas y Naturales. Universidad Nacional de Córdoba (FCEFyN – UNC). X5016GCA Córdoba. Argentina; Instituto de Investigaciones Biológicas y Tecnológicas. Consejo Nacional de Investigaciones Científicas y Técnicas (IIByT – CONICET, UNC). X5016GCA Córdoba. Argentina; Department of Structural and Chemical Biology, Centro de Investigaciones Biológicas Margarita Salas (CSIC). 28040 Madrid. Spain

## Abstract

In mammals, sperm acquire fertilization ability after a series of physiological and biochemical changes, collectively known as capacitation, that occur inside the female reproductive tract. In addition to other requirements, sperm bioenergetic metabolism has been identified as fundamental component in the acquisition of the capacitated status. Mammalian sperm produce ATP by means of two main metabolic processes, oxidative phosphorylation (OXPHOS) and aerobic glycolysis, that are localized in two different flagellar compartments, midpiece and principal piece, respectively. In mouse sperm, the occurrence of many events associated to capacitation depends on the activity of these two energy-producing pathways, leading to the hypothesis that some of these events may impose changes in sperm energetic demands. In the present study, we used extracellular flux analysis to evaluate the changes in the glycolytic and respiratory parameters of murine sperm that occur as a consequence of capacitation. Furthermore, we examined whether these variations affect sperm ATP sustainability. Our results show that capacitation promotes a shift in the usage ratio of the two main metabolic pathways, from oxidative to glycolytic. However, this metabolic rewiring does not seem to affect the rate at which the sperm consume ATP. We conclude that the probable function of the metabolic switch is to increase the ATP supply in the distal flagellar regions, thus sustaining the energetic demands that arise from capacitation.

## Introduction

After ejaculation, mammalian sperm must spend a species-specific amount of time in the female reproductive tract in order to acquire the ability to fertilize the ova. During this period, sperm undergo a series of biochemical, structural, physiological and behavioral changes that have been collectively termed capacitation [1–3]. As a consequence of capacitation, sperm become able to orient their movement in response to chemical cues originated in the cumulus-oocyte complex (chemotaxis) [4,5], and undergo the acrosomal reaction, which allows them to penetrate the oocyte vestments and fuse with the oolema [6–8]. In addition to these modifications, capacitation is also associated with a change in sperm movement pattern, termed “hyperactivation” [9]. Hyperactivated movement is characterized by highly vigorous, asymmetric flagellar beating with increased amplitude [9–12], that results in the loss of linear movement in aqueous media but increases progression efficiency in viscous media [13].

At the molecular level, sperm capacitation constitutes a highly complex process that includes modifications in the majority of cell components [14]. Capacitation involves hyperpolarization [15,16] and alterations in the architecture [17–20] and composition of the plasma membrane [19,21,22], increase in the intracellular Ca^2+^ concentration [23,24], and elevation of intracellular pH [15]. This process also involves the activation of a soluble adenylyl cyclase and the subsequent increase in the levels of cAMP [16,25–27], which in turn activates the cAMP dependent protein kinase A (pKA) pathway [16,28,29], that leads to a substantial increase in the phosphorylation of tyrosine and serine-threonine residues throughout the cell [16,30–33].

Previous studies on *in vitro* sperm capacitation have identified a series of medium components that are key for the process to take place (see [14,16,32] for reviews on the subject). Ions such as HCO^3-^ [19,34–37] and Ca^2+^ [38–40] are necessary for the activation of the phosphorylation pathways, and a cholesterol acceptor (usually serum albumin) to facilitate the efflux of cholesterol from the membrane [36]. More recently, sperm bioenergetic metabolism has been identified as an additional central component in the acquisition of the capacitated status since a number of studies have identified the activity of particular metabolic pathways as necessary for sperm capacitation (see review in [41]).

In order to sustain flagellar motility, mammalian sperm produce ATP by means of two main metabolic pathways, oxidative phosphorylation (OXPHOS) and aerobic glycolysis [42–44], which occur in different cellular compartments. While OXPHOS takes place in the mitochondria located in the midpiece (i.e., the section of the flagellum most proximate to the head), glycolysis takes place in the principal piece, where glycolytic enzymes are associated to the fibrous sheath that surrounds the axoneme [42–45]. In the domestic mouse, the activity of both metabolic routes is necessary to maintain vigorous motility [46–50] since sperm motility decreases to its eventual cessation under inhibition of glycolytic enzymes or absence of glycolytic substrates [49,51], and alterations of mitochondrial functions are associated to male infertility as a consequence of impaired sperm motility [47].

Metabolic requirements for sperm capacitation in mammals seem to be species-specific [41,45]. Numerous studies have revealed that an active glycolytic pathway is necessary for the normal occurrence of mouse sperm capacitation and its associated processes, including fertilization [46,48,52–58], and mutations that impair the catalytic function of key glycolytic enzymes [49,59–63] have been found to promote disruptions in sperm capacitation, hyperactivation, and fertilizing ability [59,60]. These findings support the idea that the physiological events associated to capacitation, such as active ion transport, activation of multiple intracellular signaling pathways, extensive protein phosphorylation, and the vigorous flagellar beating typical of hyperactivation may impose changes in sperm energetic demand [58,64].

A previous study from our laboratory [65] that examined the metabolism of sperm in the domestic mouse and other closely related species in survival (i.e., non-capacitating) conditions by means of extracellular flux analysis (Agilent-Seahorse XF24), revealed that these cells engaged in glycolysis and OXPHOS when glucose was the only substrate in the medium, and that the ratio of usage of each metabolic pathway was species-specific. However, until recently, there was little to no evidence regarding the changes in metabolic rate experienced by mouse sperm during the capacitation process. Two recent studies revealed that mouse sperm increase their glucose uptake rate when exposed to capacitating medium [66] or during chemically induced capacitation [67], in response to elevated intracellular cAMP and Ca^2+^ concentrations [66]. Furthermore, they showed the elevation of substrate uptake was associated to its utilization to fuel the increased activities of both glycolytic and respiratory pathways [67]. Nonetheless, these results only allowed for a limited analysis of the metabolic processes involved in the change from non-capacitated to capacitated, since (a) capacitation was achieved via chemical stimulation with db-cAMP and IBMX in the absence of NaHCO_3_, and (b) basal respiration and glycolysis rates were not calculated by using combinations of pathway inhibitors and, thus, phenomena such as non-mitochondrial oxygen consumption, proton leakage rate, and non-glycolytic acidification were not contemplated. Finally, there is evidence showing that an increase in mitochondrial membrane potential, that occurs only in capacitated sperm [68,69], is necessary for the acquisition of the hyperactivated movement pattern. In all, these evidences support notion that the bioenergetics of murine sperm capacitation involves changes in both, glycolytic and respiratory pathways.

The majority of studies on the relation between sperm bioenergetics and capacitation have focused on the variations that occur in ATP production mechanisms. It is intuitive to think that higher cellular ATP content would be the immediate consequence of higher ATP production rates. Nonetheless, in order to maintain cellular homeostasis and engage in any metabolic process (capacitation included), there is a need for an equilibrium between ATP synthesis and hydrolysis. Thus, the ratio between ATP production and consumption determines the level of metabolic intensity sperm may achieve and its ability to maintain adequate motility. Comparative studies in muroid rodent showed that the species with high sperm metabolic rates (ATP production) [65] are able to sustain higher ATP consumption rates [70], and consequently, faster swimming velocities for a longer time [71]. On the other hand, sperm from three *Mus* species have been shown to experience a decline in ATP content when incubated under capacitating conditions [72], suggesting that the events involved in capacitation may impose ATP demands that even the enhanced metabolic rate of capacitated sperm might not be able to sustain for a long time.

In the present study, we used extracellular flux analysis (hereafter EFA) to evaluate which changes in the glycolytic and respiratory parameters of murine sperm occur as a consequence of capacitation. Furthermore, we examined the variations in ATP sustainability that may be imposed by capacitation. We show that capacitation promotes a shift in the usage ratio of the main metabolic pathways, from oxidative to glycolytic, probably as a manner to maintain local ATP concentration in the distal flagellar regions.

## Materials and Methods

### Reagents

Unless stated otherwise, reagents were purchased from Merck (Madrid, Spain).

### Animals, sperm collection and incubation

Hybrid B6D2F1 adult male mice (3-4 months old) were purchased from ENVIGO (Barcelona, Spain). These hybrids were generated by crossing males and females from two laboratory inbred strains (DBA/2 and C57Bl/6 respectively). The animals were kept in the facilities of the Museo Nacional de Ciencias Naturales (Madrid) in individual cages under controlled temperature (20-24 °C) on a 14/10 h light/darkness photoperiod, and supplied with water and food *ad libitum*.

Care and maintenance of the mice used in this study was carried out according to the Royal Decree on Protection of Experimental Animals RD53/2013 and the European Union Regulation 2010/63, and with the approval of CSIC’s ethics committee and the Comunidad de Madrid (28079-47-A). Mice were sacrificed by cervical dislocation, which is considered as a humane sacrifice method by Spanish and European regulations. No procedures other than sacrifice for the collection of gametes were performed.

After sacrifice, mice were dissected and their caudae epididymides excised. The blood vessels, fat and surrounding connective tissue was removed from the caudae. Each cauda epididymis was placed in a 35 mm Petri dish containing 0.4 ml of one of two variants of culture medium pre-warmed to 37 °C. One cauda epididymis was placed in non-capacitating medium under air and the other one was placed in capacitating medium under 5% CO_2_/air. The compositions of both media were based on a Hepes-buffered modified Tyrode’s medium [35], supplemented with albumin, lactate and pyruvate (pH = 7.4, osmolality = 295 mOsm kg^−1^). The composition of the non-capacitating medium was: 132 mM NaCl, 2.68 mM KCl, 0.49 mM MgCl_2_.6H_2_O, 0.36 mM NaH_2_PO_4_.2H_2_O, 5.56 mM glucose, 20 mM HEPES, 1.80 mM CaCl_2_, 0.02 mM phenol red, 0.09 mM kanamycin, 4 mg ml^−1^ fatty acid-free BSA, 20 mM Na lactate, and 0.5 mM Na pyruvate. The capacitating medium differed from the non-capacitated medium in that 15 mM NaHCO_3_ was added and NaCl adjusted to maintain the osmolality.

Three to five incisions were performed in the distal region of the caudae, and sperm were allowed to swim out for 5 min [72]. For most experiments (see below), sperm from two mice were used in each experiment. For the latter, one epididymis from each of two males was placed in a petri dish (total of two per dish) and the sperm of both was mixed during swim out. After swim out, the epididymal tissue was discarded and the sperm suspension was collected into a plastic tube under a suitable atmosphere. Sperm concentrations were estimated in each sperm suspension by using a modified Neubauer chamber, and adjusted to 100 × 10^6^ sperm ml^−1^ with the corresponding medium. Concentration-adjusted sperm suspensions were then incubated during 1 h at 37 °C under suitable atmospheres since at this time, mouse sperm population tend to reach the peak of capacitated cells [72]. In all procedures, large-bore pipette tips were used to minimize mechanical damage to spermatozoa.

In the case of sperm ATP consumption measurements, only one male was used in each experiment. Thus, sperm from one cauda epididymis was collected in each medium. In this case, the sperm concentration of each suspension was adjusted to 20 × 10^6^ sperm ml^−1^ prior to the 1-hour incubation time.

### Sperm capacitation

Sperm capacitation status was assessed at the end of the 1 h incubation period, and prior to extracellular flux analysis. Capacitation was assessed using chlortetracycline (CTC, C4881) combined with a vital stain [72,73]. Briefly, after a 1 h incubation under different conditions, 100 μl of the sperm suspension were mixed with 50 μl of 6 μg/ml Hoechst 33258 bisbenzamide (B2883) and incubated during 1 min in the dark. Subsequently, the samples were centrifuged for 2 min at 100 ×g, the supernatant was discarded, the pellet resuspended in 100 μl of medium and fixed with an equal volume of 2% glutaraldehyde-0.165 M sodium cacodylate solution.

Immediately before evaluation, an aliquot of 20 μl of sperm suspension was added to 20 μl of 250 μM CTC solution and incubated in the dark for 3 min. Spermatozoa were observed at 1000× magnification under fluorescence and phase contrast microscopy simultaneously (E-600 microscope, Nikon). Pre-fixation viability of the spermatozoa was assessed using a Nikon UV-2A 330-nm filter and fluorescence emission via a DM 400 dichroic mirror. Only the cells that excluded the Hoechst stain were considered viable. CTC staining patterns were observed using a Nikon BV-2A 405-nm filter and fluorescence emission with a DM 455 dichroic mirror, and the following patterns were distinguished [72]: (a) F (non-capacitated sperm): the head of spermatozoa was uniformly stained with CTC; (b) B (capacitated sperm): the post-acrosomal region of the head was not stained with CTC; and (c) AR (without the sperm acrosome): the head of the sperm cell showed little or no CTC staining. Capacitation status was evaluated in 100 viable spermatozoa with intact acrosomes per sample and expressed as the percentage of sperm showing staining pattern “B”.

### Assessment of sperm metabolic rates

An XFp extracellular flux analyzer (Agilent Seahorse, Santa Clara, CA) was used to measure oxygen consumption rate (OCR) and extracellular acidification rate (ECAR) in real time. In this system, OCR indicates the level of respiratory activity in a population of cells, and ECAR, an estimate of the rate of lactate release to the medium, is used as a proxy for aerobic glycolytic activity. After 1 h incubation in differential conditions (i.e., capacitation period), the cells were transferred to an 8-well XFp plastic microplate. As a substantial difference from the original method developed by Tourmente et al. [65], sperm were not attached to the bottom of the well. Thus, measurements were obtained from a free-swimming sperm population. This required an increase the number of sperm added to each well in order to ensure that the 2 μl closed micro-chamber, formed between the sensor and the bottom of the plate when the instrument is measuring, contained enough cells to detect a reliable signal. Three wells were seeded with 80 μl of sperm suspension (approximately 8 x 10^6^ sperm) incubated under non-capacitating conditions, other three wells were seeded with sperm suspension incubated under capacitating conditions, and two wells were left without cells to perform background corrections. The plate was centrifuged for 1 min at 1300 xg in each direction to temporarily set the cells at the bottom of the well. The supernatant was removed from each well and 200 μl of unbuffered registry medium were added to the 8 wells of the plate, thus resuspending the cells. Since the presence of a pH buffer in the medium would impede an accurate measurement of ECAR, an unbuffered registry medium was used for the measurements and dilution of the metabolic modulators. In this medium, Hepes and NaHCO_3_ were replaced by NaCl to preserve osmolality, and pH adjusted to 7.4 at 37 °C. After medium replacement, the plate with the sperm was placed in the XFp, and a 12 min equilibration step was allowed before measurements.

OCR and ECAR were measured at 37 °C and the measurement cycle consisted of 2 min of mixing, 1 min of waiting, 3 min of measuring. In a first set of experiments (n = 4), we assessed whether stable measurements of sperm metabolism could be obtained from mouse sperm samples before the addition of any compound. Thus, OCR and ECAR were recorded for 48 min (8 cycles). Subsequently, 1 μM rotenone and 1μM antimycin A (inhibitors of the mitochondrial respiratory chain complexes I and III respectively) were added to the wells, and both metabolic variables were monitored for an additional 12 min (10 cycles, 60 min). Oxygen levels showed a marked descent during the measurement phase that fully recovered during the mixing and waiting phases (Fig. S1), indicating that the sperm concentration used in the experiment did not lead to hypoxia.

For the second set of experiments, we measured sperm metabolism both before and after the addition of metabolically active compounds, in order to calculate a range of metabolic parameters. OCR and ECAR were measured for 36 min (6 cycles) to establish a baseline value (hereafter “basal”). Subsequently, one of three metabolic modulators was added to the wells, and sperm metabolism was recorded for 3 additional cycles (18 min). Metabolic modulators were: (a) 5 μM oligomycin A, an inhibitor of the mitochondrial ATP synthase, (b) 1 μM carbonyl cyanide p-trifluoro-methoxyphenylhydrazone (FCCP), an uncoupler of mitochondrial OXPHOS, and (c) 50 mM 2-deoxy-d-glucose (2DOG), a glucose analog that competitively inhibits the first step of glycolysis. Finally, 1 μM rotenone and 1μM antimycin A were added to the wells for two final measurements (11 cycles, 66 min). Oligomycin and FCCP were used in four separate experiments, while 2DOG was used in three separate experiments (total n = 11 for the second set). For all experiments, the first two measurements were discarded since, according our previous experience, these tend to show a higher degree of instability. Thus, the first OCR and ECAR values considered for this study were taken 12 min after the beginning of the measurements.

At the end of the experiment, the sperm suspension in each well was mixed by pipetting, and 10 μl collected to estimate sperm concentration using a modified Neubauer chamber. Sperm numbers in the wells were calculated and the OCR and ECAR values for each well were normalized by the number of sperm present in the 2 μl measurement volume (reported as amol of O_2_ min^−1^ sperm^−1^, and nano-pH min^−1^ sperm^−1^, respectively).

### Calculation of metabolic parameters

The addition of the metabolic modulators used in the second set of EFAs allowed us to calculate several metabolic parameters (Table S1) that have more informative value than the raw normalized measurements. They also allowed for a more accurate comparison between the two incubation conditions [74,75]. Metabolic parameters were calculated for each well as the average value of the measurements taken for that well in condition A minus the average value of the measurements taken for that well in condition B (see Table S1 for a description of the conditions and Fig. S2 for a graphical representation). Values corresponding the basal state (no additions) were averaged across measurement cycles 3-6, values corresponding to oligomycin and 2DOG additions were averaged across cycles 7-9, and values corresponding to A+R additions were averaged across cycles 10-11. Exceptionally, for the calculation of the stimulating effects of FCCP and Oligomycin on OCR and ECAR respectively, the highest value among cycles 7-9 was used instead of the average.

In order to facilitate comparisons with previous and future studies, we also calculated the response of OCR and ECAR to metabolic modulators as a percentage of the averaged baseline values (measures 3-6) for each well. In the case of OCR, both the baseline and the stimulated/inhibited values were first corrected by subtracting the average of the measures after the addition of A+R (cycles 10-11).

### Sperm ATP content

A separate set of experiments (n = 5), was performed in order to compare sperm ATP consumption rates between sperm incubated in the different conditions. After 50 min of incubation under capacitating or non-capacitating conditions, each sperm suspension was divided in two aliquots. The first sperm aliquot (hereafter “control”) was used to monitor basal ATP content throughout the consumption experiment. Sperm ATP content was assessed in this aliquot at 55 and 70 min of incubation. Inhibitors of OXPHOS (1 μM antimycin A, 1 μM rotenone, and 5 μM oligomycin) and glycolysis (50 mM 2DOG) were diluted in the corresponding medium and added to the second aliquot (hereafter “inhibition” treatment) after 60 min of incubation in order to make sperm able to consume ATP without producing it [70]. Sperm ATP content was assessed in the inhibition treatment immediately after the addition of inhibitors and after 2, 4, 6, 8, and 10 min. Although the addition of rotenone and antimycin A would suffice to stop OXPHOS-derived ATP production, oligomycin was used to an artefactual ATP depletion due to the reverse action of the mitochondrial ATP synthase [55]. Since OXPHOS inhibitors were dissolved in DMSO prior to the addition to the sperm suspension, this solvent was added to the control treatment prior to its first ATP content assessment to an equivalent concentration (0.7% v/v).

Sperm ATP content was measured using a luciferase-based ATP bioluminescent assay kit (Roche, ATP Bioluminescence Assay Kit HS II) based on the protocol of Tourmente et al. [70]. Sperm suspension was diluted 1:10 in the corresponding medium, and a 100 μl aliquot of diluted sperm suspension was mixed with 100 μl of Cell Lysis Reagent and frozen in liquid N_2_. At the end of the experiment, all the aliquots were thawed at room temperature for 5 min. The resulting cell lysate was centrifuged at 12,000 × g for 2 min, and the supernatant was recovered and frozen in liquid N_2_. Bioluminescence was measured in triplicate in 96-well plates using a luminometer (Biotek Synergy, Biotek Instruments Inc.). Auto injectors were used to add 50 μl of luciferase reagent to 50 μl of sample in each well, and, following a 1 s delay, light emission was measured over a 5 s integration period. Standard curves were constructed using ATP standards diluted with the corresponding media and Cell Lysis Reagent in a proportion equivalent to that of the samples. ATP content was expressed as amol sperm^−1^.

### Statistical analyses

All analyses were conducted using R version 4.1.3 (R Foundation for Statistical Computing, Vienna, Austria), with *α* = 0.05. To assess whether incubation conditions affected the percentage of capacitated sperm, we applied a generalized linear mixed effect model (GLMM, *mixed* function, *afex* package) with binomial distributions and “logit” link function, including incubation conditions as a fixed factor with 2 levels (non-capacitating vs. capacitating), and experiment as random factor. In the first set of EFAs the stability of the OCR and ECAR measures during the assay was assessed via linear mixed effect models (LMM, *mixed* function, *afex* package) using time as a fixed factor with 8 levels (one per measurement cycle), and experiment as random factor. In the second set of EFAs, we analyzed the effect of incubation conditions and metabolic modulators on OCR and ECAR expressed as percentages relative to the baseline using LMMs with incubation medium and treatment (basal, oligomycin, FCCP, 2DOG, and A+R) as fixed factors, and experiment as random factor. The inclusion of the experiment as a random factor allowed for the consideration of each well as an individual value in the statistical analyses, thus decreasing the impact of outliers and increasing the statistical powers of the tests by controlling for the between-experiment variability [76]. The values of the metabolic parameters calculated after EFAs were compared between incubation conditions using LMMs with incubation medium as a fixed factor and experiment as a random factor.

Non-linear regression models (*ngls* function, *nlme* package) were used to estimate sperm ATP consumption rates for sperm incubated in capacitating and non-capacitating conditions and subsequently treated with inhibitors. First, a general regression curve (ATP = *a* x *e*^*b* x time^) based on the values of ATP content at the six sampling time points was estimated for each condition using the pooled measurements of the four experiments. The parameter “b” was defined as the ATP consumption rate, that is, the exponential rate of decline in ATP content as a function of time. In addition, a second ATP consumption rate was estimated for each individual sample and the “b” exponents of the curves were compared between conditions by means of a LMM with experiment as random factor.

All variables were log_10_-transformed for statistical purposes in all analyses with the exception of the values expressed as percentages, and the estimated ATP consumption rates. Significant differences between levels of fixed factors were analyzed via *post-hoc* estimated marginal means tests (*pairwise* function of the *emmeans* package).

## Results

After 1 h of incubation post collection, approximately 50 % of the live sperm incubated in capacitating conditions presented the pattern “B” of CTC staining. On the other hand, only 13 % of the live sperm showed such pattern when incubated under non-capacitating conditions, confirming that incubation in capacitating medium promoted a significant increase in the percentage of capacitated sperm in the samples (mean ± SE: 51.5 ± 1.59 vs 13.17 ± 1.59; GLMM: *X^2^* = 428.52, *p* < 0.001; Fig. 1).

**Figure 1.**
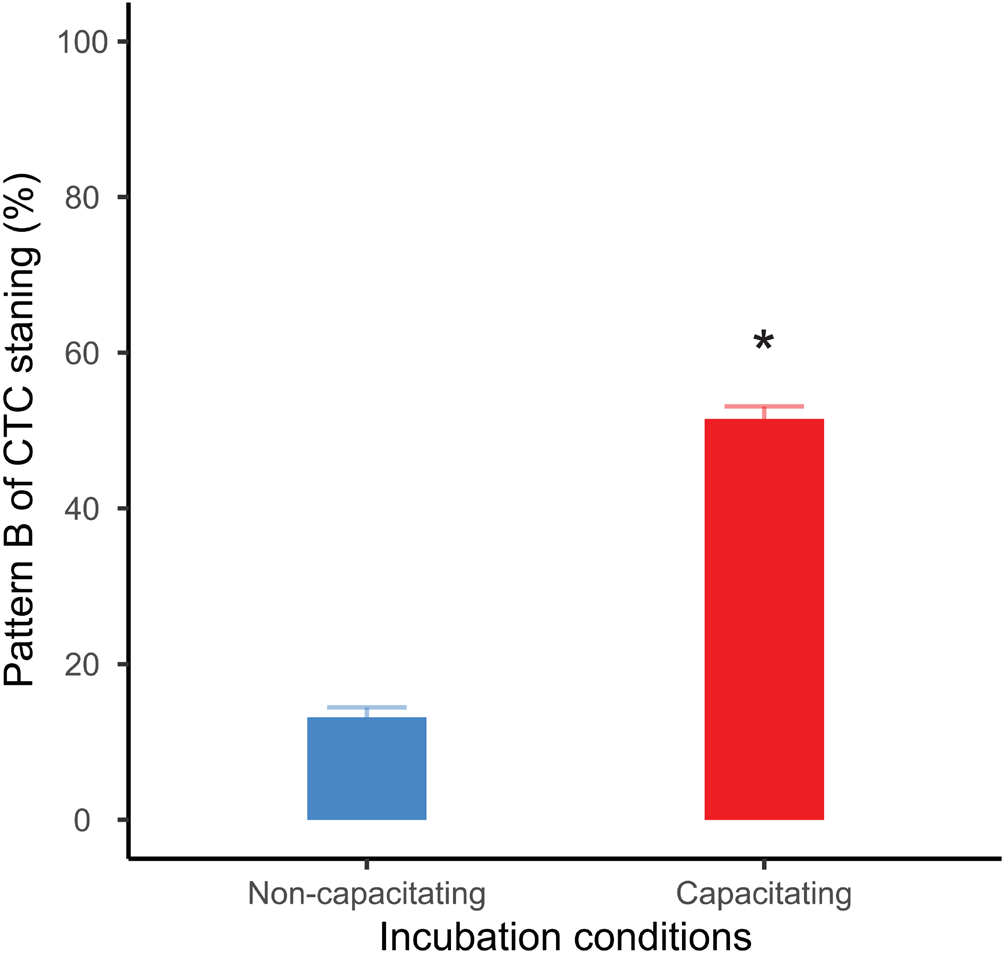
Percentage of live capacitated sperm (CTC pattern “B”) after one hour of incubation. Bars correspond to means + standard error. Blue bar: sperm incubated in non-capacitating conditions; red bar: sperm incubated in capacitating conditions. Asterisk indicates significant differences (*p* < 0.05) in percentages between incubation media.

The first set of EFAs showed that experimental time had no significant effects on OCR values before A+R addition in both incubation conditions (non-capacitating: LMM *X^2^* = 4.13, *p* = 0.531; capacitating: LMM *X^2^* = 2.56, *p* = 0.768); Fig. S3A, C). Similarly, experimental time did not have a significant effect on ECAR for any of the two incubation conditions (non-capacitating: LMM *X^2^* = 1.59, *p* = 0.903; capacitating: LMM *X^2^* = 8.66, *p* = 0.123; Fig. S3 B, D). These results indicate that the modification applied to the EFA technique allowed us to obtain stable OCR and ECAR values for at least 54 minutes after introducing the samples into the XFp analyzer.

The OCR and ECAR profiles for the second set of EFAs in both incubation conditions are shown in Fig. 2. In both non-capacitating and capacitating sperm, cells responded as expected to the injection of metabolically active compounds (Figs. 2, Table 1). Thus, OCR significantly increased when exposed to the uncoupler FCCP, decreased when exposed to the ATP synthase inhibitor oligomycin, and further diminished when exposed to antimycin and rotenone (Fig. 2A, B, Table 1). Furthermore, sperm OCR was reduced when exposed to the glycolytic inhibitor 2DOG. ECAR values decreased as expected in response to the addition of 2DOG in both incubation conditions, and increased when respiration was inhibited by the addition of oligomycin (Fig. 2C, D, Table 1).

**Figure 2.**
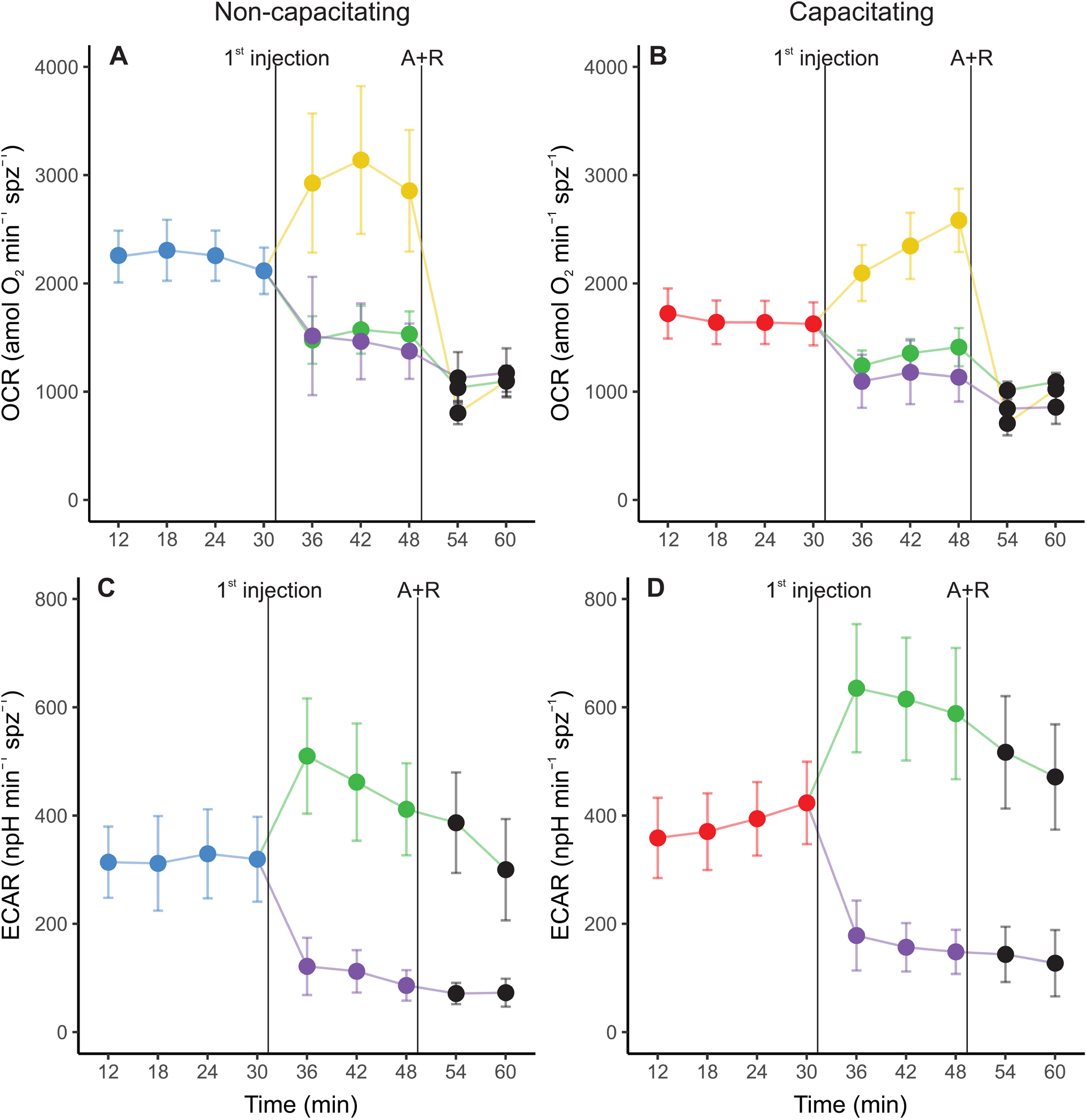
Real-time measurements of oxygen consumption rate (OCR) (A, C) and extracellular acidification rate (ECAR) (B, D) in mouse sperm before and after treatment with metabolic modulators. Sperm were incubated during 1 hour in non-capacitating (A, C) and capacitating (B, D) conditions prior to extracellular flux analysis. Values have been normalized by sperm numbers inside each well. Symbols and whiskers correspond to means ± standard error. Time = 0 was defined as the start of the first measurement cycle; measurement cycles 3 to 11 are reported. The line labeled as “1st injection” marks the addition of either 5 μM oligomycin, 1 μM FCCP, or 50 mM 2DOG; the line labeled “A+R” marks the addition of 1 μM antimycin A + 1 μM rotenone. Blue and red symbols: values before any addition (basal state) in non-capacitating and capacitating conditions respectively; green symbols: after oligomycin injection; yellow symbols: after FFCP injection; purple symbols: after 2DOG injection; black symbols: after A+R injection.

**Table 1.** Effect of incubation conditions and metabolic modulators on mouse sperm OCR and ECAR.

| Dependent variable | Independent variable | $\chi^2$ | $p$ |
| --- | --- | --- | --- |
| OCR (%) | Incubation medium | 44.07 | **<0.001** |
|  | Treatment | 673.72 | **<0.001** |
|  | Interaction | 113.44 | **<0.001** |
| ECAR (%) | Incubation medium | 16.71 | **<0.001** |
|  | Treatment | 325.89 | **<0.001** |
|  | Interaction | 34.65 | **<0.001** |

Independent variables were tested as percentages relative to the basal state. $\chi^2$ and $p$ values were estimated using LMMs and likelihood ratio tests. Incubation medium (non-capacitating or capacitating) and treatment (basal, 5 μM oligomycin, 1 μM FCCP, 50 mM 2DOG, and 1 μM antimycin + 1 μM rotenone) were used as fixed factors and experiment as random factor. Significant differences ( $p < 0.05$ ) are shown in boldface.

Sperm populations incubated under both conditions showed qualitatively similar responses to metabolic modulators when OCR and ECAR were expressed as percentages relative to the basal state. Exposure to oligomycin caused a reduction of approximately 71 % in OCR (Fig. 3A, Fig. 4A, Table 1). The addition of 2DOG promoted a decline in OCR that was similar to that produced by exposure to oligomycin (^~^ 70%) (Fig. 3C, Fig. 4A, Table 1) and a decrease of approximately 60% in ECAR (Fig. 3E, Fig. 4B, Table 1). On the other hand, some treatments elicited different responses in non-capacitating and capacitating sperm. The increase in OCR associated to the addition of FCCP was significantly higher in sperm incubated under capacitating conditions (319 % vs 192 %) (Fig. 3B, Fig. 4A, Table 1). Also, the rise in ECAR as a response to oligomycin in sperm incubated under capacitating conditions surpassed that of sperm incubated in non-capacitating conditions (90 % vs 60 %) (Fig. 3D, Fig. 4B, Table 1).

**Figure 3.**
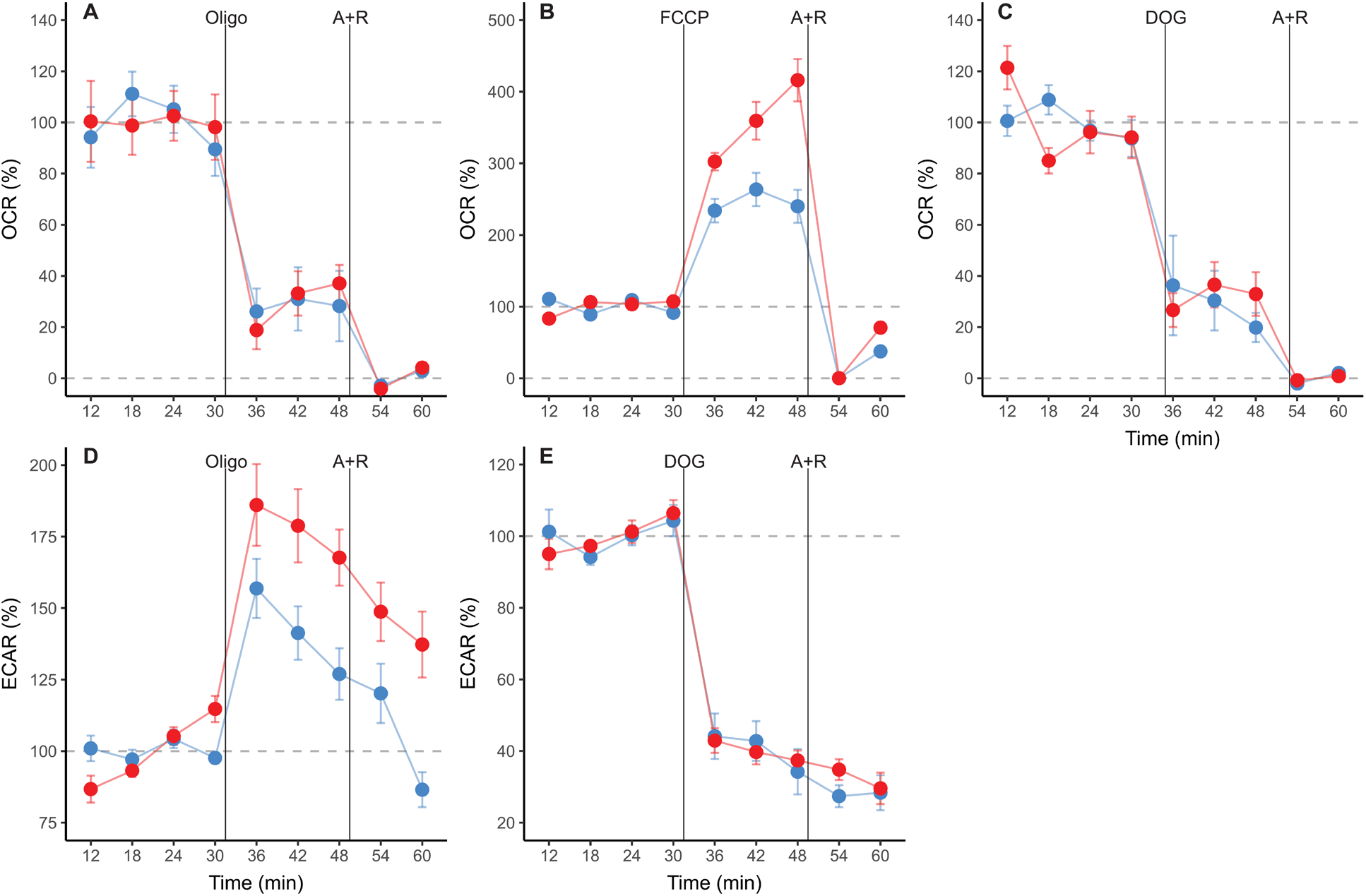
Real-time measurements of oxygen consumption rate (OCR) (A, B, C), and extracellular acidification rate (ECAR) (D, E) in mouse sperm before and after treatment with metabolic modulators. Values are expressed as percentages relative to the basal state (100%), which was defined as the mean of the four measurement cycles before any addition and is represented by the horizontal dashed line. Sperm were incubated during 1 hour in non-capacitating (blue symbols) and capacitating (red symbols) conditions prior to extracellular flux analysis. Symbols and whiskers correspond to means ± standard error. Time = 0 was defined as the start of the first measurement cycle; measurement cycles 3 to 11 are reported. Dashed grey lines indicate 100 % and 0% levels. The first vertical line marks the addition of either 5 μM oligomycin, 1 μM FCCP, or 50 mM 2DOG; the line labeled “A+R” marks the addition of 1 μM antimycin A + 1 μM rotenone.

**Figure 4.**
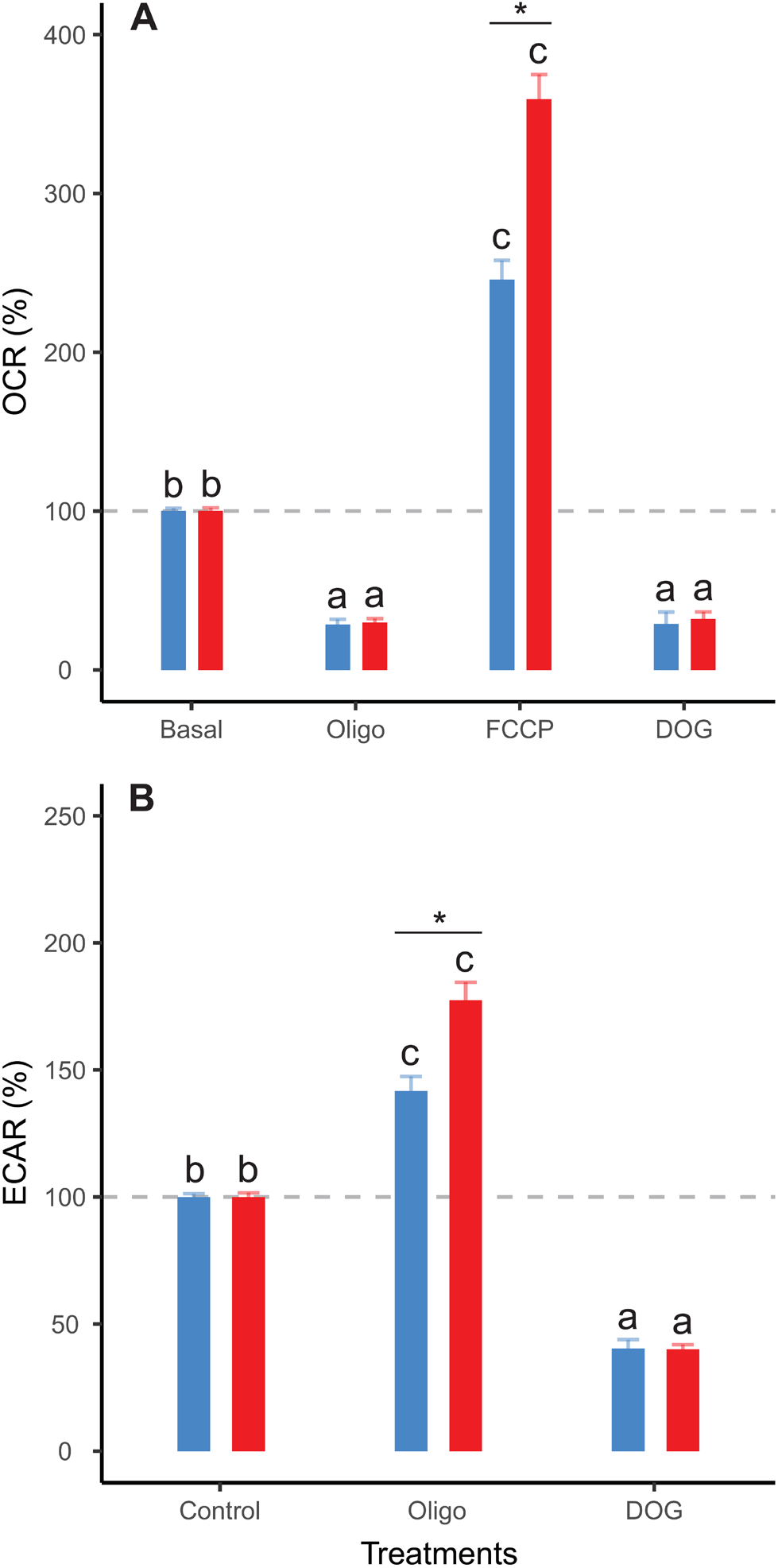
Effect of metabolic modulators on mouse sperm OCR (A), ECAR (B) expressed as percentages relative to the basal state. Sperm were incubated in non-capacitating (blue bars) and capacitating (red bars) conditions prior to extracellular flux analysis. Bars correspond to means + standard error. Treatments represent sperm metabolic rates before any addition (Basal) or after the addition of 5 μM oligomycin (Oligo); 1 μM FCCP (FCCP); 50 mM 2DOG (DOG); or 1 μM antimycin A + 1 μM rotenone (A+R). Dashed grey lines indicate 100 % level. Different letters indicate significant differences (*p* < 0.05) between treatments for the same incubation condition in a *post-hoc* marginal means test. Asterisks indicate significant differences (*p* < 0.05) between incubation conditions for the same treatment in a *post-hoc* marginal means test.

Clear differences between incubation conditions emerged when the OCR and ECAR measurements were used to calculate and compare a range of metabolic parameters (Table S1). Sperm incubated under non-capacitating conditions presented higher values in nearly all parameters associated with respiratory metabolism (basal respiration rate, proton leak, respiratory ATP production, and maximum respiration rate) (Fig. 5A-D, Table 2), with the exception of spare respiratory capacity, which did not show any significant differences between incubation conditions (Fig 5E, Table 2). Conversely, the two parameters associated with aerobic glycolysis (basal glycolysis rate and glycolytic reserve) showed higher values in sperm incubated under capacitating conditions (Fig. 5G, H, Table 2). The OCR/ECAR ratio supported this trend, since it was significantly higher in the sperm incubated in non-capacitated conditions (Fig. 5F, Table 2). In all, these results confirm the occurrence of a switch in the usage ratio of the two energy supply pathways as a consequence of sperm capacitation.

**Figure 5.**
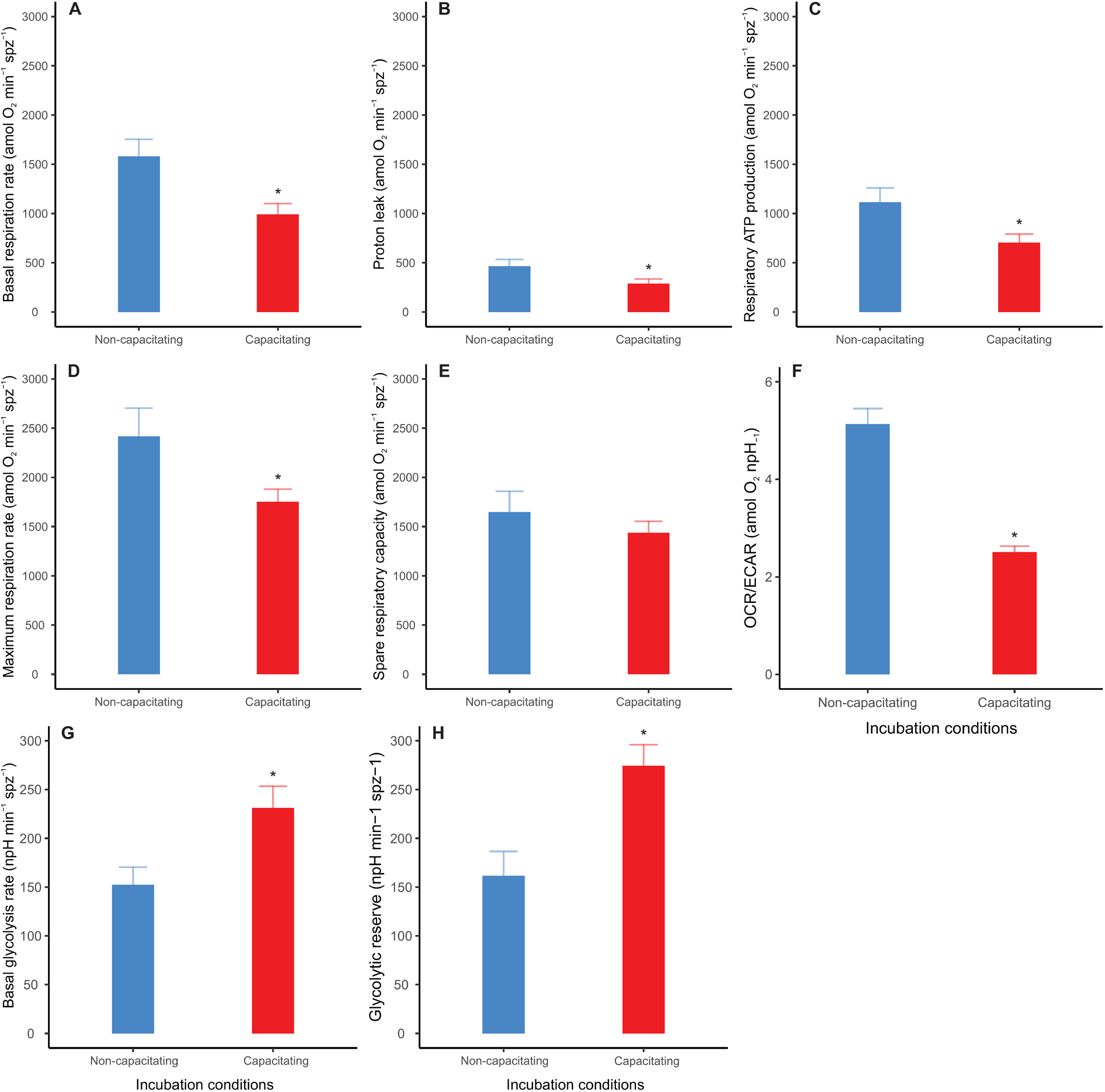
Effect of incubation conditions on mouse sperm metabolic parameters and OCR/ECAR ratio. OCR/ECAR ratios correspond to basal state (before any additions). Sperm were incubated in non-capacitating (blue bars) and capacitating (red bars) conditions prior to extracellular flux analysis. Bars correspond to means + standard error. Asterisk indicates significant differences (*p* < 0.05) in values between incubation media.

**Table 2.** Effect of incubation conditions on mouse sperm metabolic parameters and OCR/ECAR ratio.

| Dependent variable | Non-capacitating | Capacitating | $\chi^2$ | $p$ |
| --- | --- | --- | --- | --- |
| Basal respiration rate (amol O<sub>2</sub> min<sup>-1</sup> spz<sup>-1</sup>) | 1578.99 ± 174.46 | 991.53 ± 109.53 | 10.15 | **0.001** |
| Proton leak (amol O<sub>2</sub> min<sup>-1</sup> spz<sup>-1</sup>) | 464.74 ± 68.79 | 287.08 ± 102.28 | 5.55 | **0.018** |
| Respiratory ATP production (amol O<sub>2</sub> min<sup>-1</sup> spz<sup>-1</sup>) | 1114.25 ± 145.73 | 704.45 ± 86.32 | 6.28 | **0.012** |
| Maximum respiration rate (amol O<sub>2</sub> min<sup>-1</sup> spz<sup>-1</sup>) | 2416.82 ± 286.37 | 1750.46 ± 129.81 | 5.37 | **0.021** |
| Spare respiratory capacity (amol O<sub>2</sub> min<sup>-1</sup> spz<sup>-1</sup>) | 1647.48 ± 211.50 | 1437.26 ± 115.76 | 0.90 | 0.342 |
| Basal glycolysis rate (npH min<sup>-1</sup> spz<sup>-1</sup>) | 152.25 ± 18.15 | 231.10 ± 22.31 | 8.53 | **0.003** |
| Glycolytic reserve (npH min<sup>-1</sup> spz<sup>-1</sup>) | 161.67 ± 24.89 | 274.34 ± 21.79 | 10.16 | **<0.001** |
| OCR/ECAR (amol O<sub>2</sub> npH<sup>-1</sup>) | 5.13 ± 0.32 | 2.51 ± 0.12 | 102.02 | **<0.001** |

Values shown mean ± standard error for each incubation condition. OCR/ECAR values correspond to basal state. $\chi^2$ and $p$ values were estimated using LMMs and likelihood ratio tests. Incubation condition (non-capacitating or capacitating) was used as fixed factor and experiment as random factor. Significant differences ( $p < 0.05$ ) are shown in boldface.

Incubation conditions did not produce significant variations in the ATP content of sperm that had not been treated with metabolic inhibitors (LMM *X^2^* = 0.45, *p* = 0.503). However, these cells showed a slight (^~^20%) but significant reduction in ATP content throughout the experimental time (Fig. 6A) (LMM *X^2^* = 5.60, *p* = 0.018). Conversely, sperm incubated in the presence of inhibitors showed a severe and sustained decrease in ATP content over time in both incubation conditions (Fig. 7A). Non-linear regression analyses performed by pooling the measurements of the five individuals yielded ATP consumption curves that significantly adjusted to the “ATP = *a* x *e*^*b* x time^” model in both incubation conditions (non-capacitating: ATP = 37.09 x *e*^-0.21 x time^, *F_b_* = 50.25, *p_b_* < 0.001; capacitating: ATP = 30.50 x *e*^-0.19 x time^, *F_b_* = 32.33, *p_b_* < 0.001) (Fig. 6B). In order to compare ATP consumption between incubation conditions, we estimated ATP consumption curves for each individual in each condition and compared the “*b*” exponents of the curve. There were no significant differences (LMM *X^2^* = 0.15, *p* = 0.712) between the values of the exponents from curves estimated for non-capacitating (*b* = −0.20 ± 0.02) and capacitating conditions (*b* = −0.19 ± 0.03).

**Figure 6.**
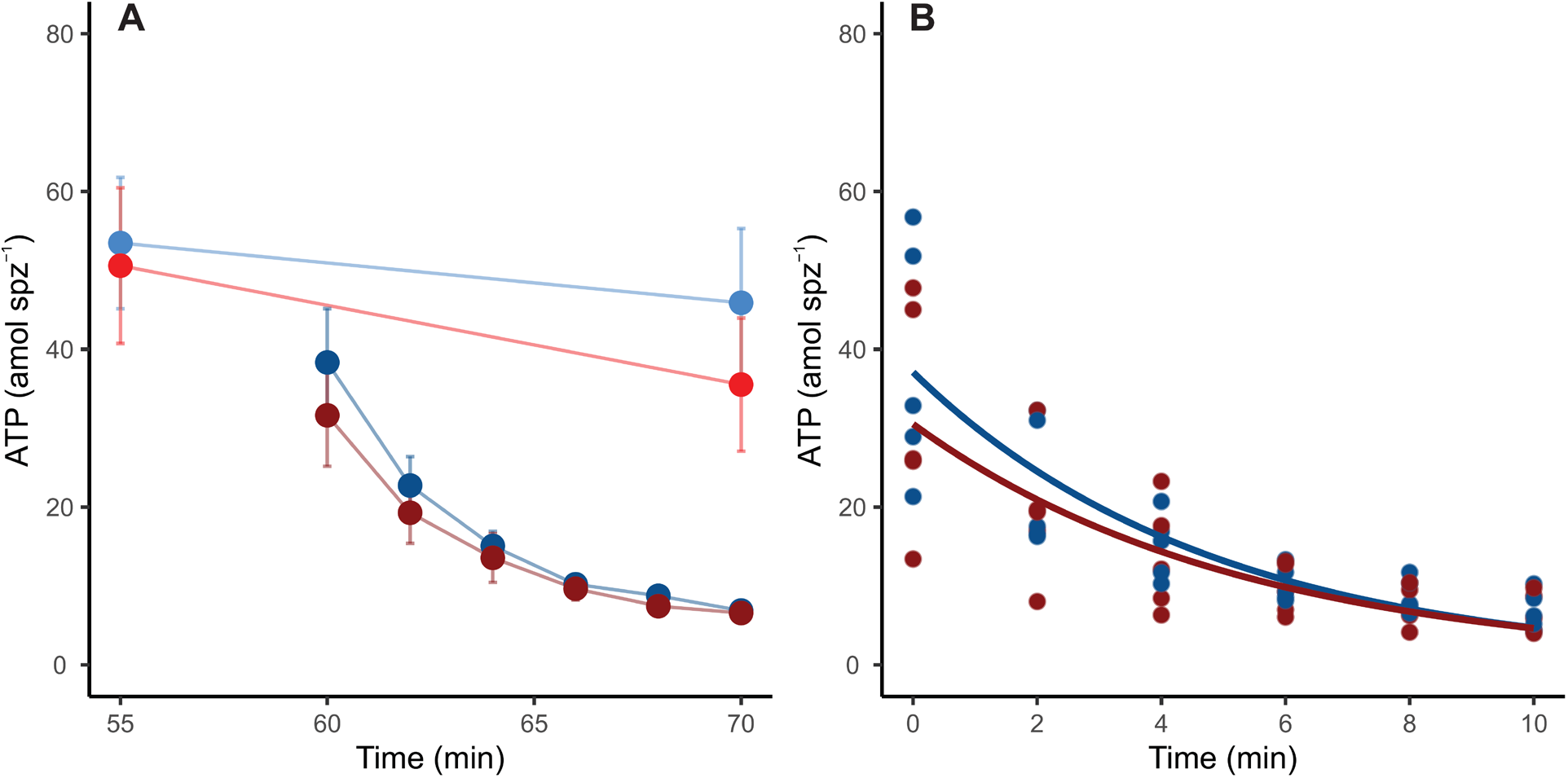
ATP consumption in mouse sperm in non-capacitating (blue symbols and lines) and capacitating (red symbols and lines) conditions. (A) ATP content in response to the addition of metabolic inhibitors to sperm in both incubation conditions. Light lines and symbols represent control conditions (no inhibitors added); dark lines and symbols represent metabolically inhibited sperm (5 μM oligomycin, 50 mM 2DOG, 1 μM antimycin A, and 1 μM rotenone were added). B) Lines represent estimated sperm ATP consumption curves in non-capacitating (ATP = 37.09 x *e*^-0.21 x time^) and capacitating (ATP = 30.50 x *e*^-0.19 x time^) conditions. Symbols represent individual measurements.

**Figure 7.**
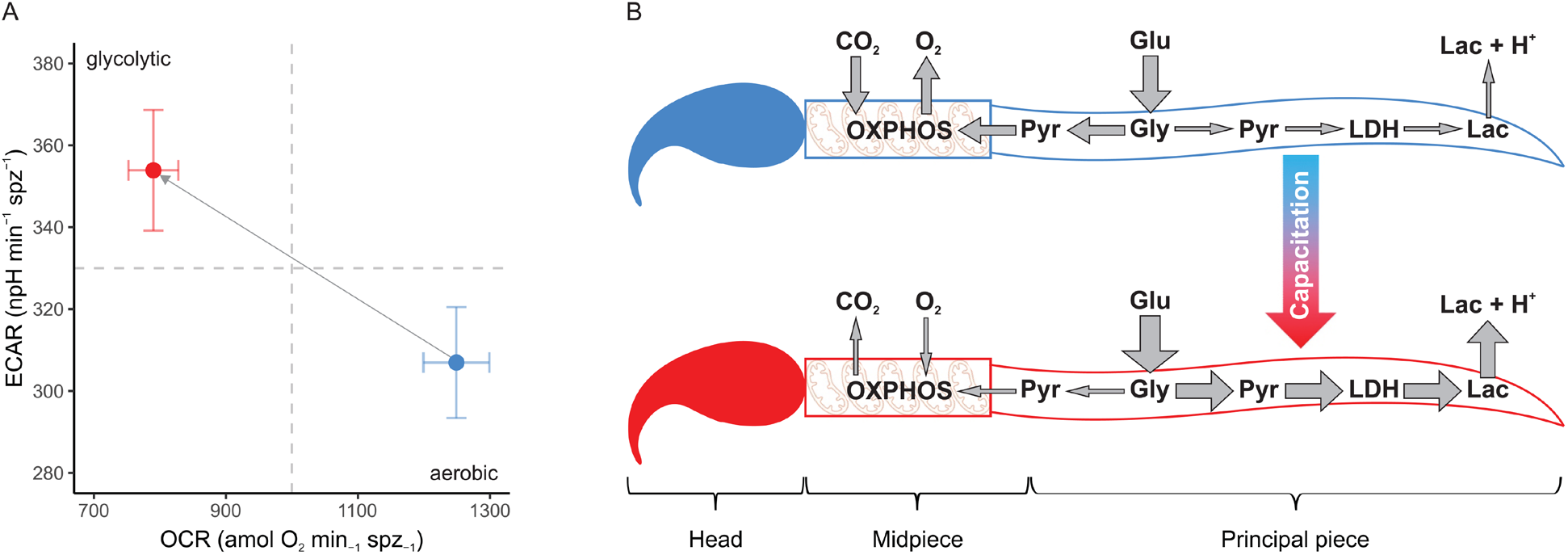
Changes in metabolic phenotype promoted by sperm capacitation in murine sperm. (A) Metabolic phenotype plot of mouse sperm depicting the usage of the two main metabolic pathways (OXPHOS and anaerobic glycolysis). Symbols and whiskers correspond to means ± standard error. Blue symbols: non-capacitating conditions. Red symbols: capacitating conditions. Dashed grey lines indicate the midpoint in the measurement range for both variables. (B) Graphical depiction of conclusions. Capacitation promotes a metabolic shift in the mouse sperm, increasing the uptake of glucose and altering the proportion of pyruvate that enters each metabolic pathway. A greater proportion of pyruvate is reduced into lactate by the enzyme lactate dehydrogenase and transported to the extracellular environment along with protons, while a relatively minor proportion is transported to the mitochondria in the midpiece, thereby entering the OXPHOS pathway. Lac: lactate. LDH: lactate dehydrogenase. Pyr: pyruvate. Gly: glycolysis. Glu: glucose.

## Discussion

The results of our study revealed that the development of capacitation in murine sperm is associated with changes in the energy metabolism of these cells. In the present study, sperm capacitation was achieved through the incubation of spermatozoa during 1 hour in a medium containing BSA, Ca^2+^ and NaHCO_3_. Sperm capacitation was confirmed by CTC staining. The incubation period and medium components required for murine sperm capacitation have been described in previous studies [30,35,72,77], and the changes in CTC staining patterns have shown to strongly correlate to markers of sperm capacitation, such as progesterone induced acrosome reaction [78] and hyperactivated movement [72]. Thus, we can confidently infer that the observed rise in CTC pattern “B” after incubation in capacitating conditions was indicative of an increased proportion of capacitated cells within the sample. Moreover, sperm populations incubated in capacitating medium are a mix of capacitated and non-capacitated cells, thus, the metabolic variations observed as a consequence of sperm capacitation are underestimated since they are attenuated by the presence of non-capacitated sperm.

In the present study, the increase in the proportion of capacitated cells in the sperm populations was associated with a change in their metabolic phenotype (Fig. 7A). Parameters associated to respiratory metabolism (basal respiration, proton leak, respiratory ATP production and maximal respiration) presented higher values in sperm populations with low percentage of capacitated cells. Conversely, sperm populations enriched in capacitated sperm relied more on glycolysis, as evidenced by their higher basal glycolysis and glycolytic reserve values. This change in metabolic profile is summarized in the OCR/ECAR ratio, which is decreases by 50% after incubation in capacitating medium. In fact, the magnitude of this metabolic switch may be underestimated by the nature of the measuring system (i.e., using ECAR as a proxy of the rate of lactate formation) since HCO3^−^ formation, derived from the production of CO_2_ via OXPHOS, will contribute to total ECAR in variable proportions depending on cell type and substrate [79]. In our study, the decrease in sperm O_2_ consumption (OXPHOS) associated with capacitation leads to a stoichiometric reduction in CO_2_ production. Thus, the increase in ECAR from lactate excretion (glycolysis), that results from the metabolic switch associated to capacitation, is partially attenuated because the CO_2_ contribution has diminished.

Sperm incubated in both media (capacitating and non-capacitating), reacted as expected to the addition of metabolic modulators. While OCR decreased in response to oligomycin, antimycin and rotenone, and increased in response to FCCP, ECAR was reduced when sperm were exposed to 2DOG. These results allowed us to validate the robustness of the technique, and to calculate, for the first time in murine sperm, a range of relevant metabolic parameters that allow for the comparison between studies and species. Importantly, this is, to the best of our knowledge, the first study to achieve robust metabolic measurements using XF EFA technology in freely moving sperm, since previous work using these techniques necessitated the use of attached cells [65,67,80]. Nonetheless, while the metabolism of mouse sperm responded similarly to metabolic modulation regardless of incubation conditions, there were quantitative differences associated with capacitation in the level of these responses. The increased elevation of glycolytic rate by capacitated sperm in response to OXPHOS inhibition (glycolytic reserve) was caused by a more pronounced reaction from a higher basal glycolytic rate. On the other hand, the proportionally higher stimulation of the respiratory chain exhibited by sperm in capacitating medium was a consequence of their decreased basal respiration rate, as evidenced by their lower maximum respiration in comparison to their non-capacitated counterparts. These evidences further support the notion of a metabolic rewiring of capacitated spermatozoa in favor of aerobic glycolysis over OXPHOS.

A recent EFA study on murine sperm [67] revealed an increase in rates of OXPHOS and aerobic glycolysis that were both concomitant with the emergence of capacitation markers, and with an increase in glucose uptake. Although, at first hand, the observations in this earlier study [67] seem contradictory with our results, there are important differences between the experimental designs of both studies that make comparison difficult. The measurements in Balbach *et al*. [67] were obtained by subjecting sperm to capacitating conditions within the extracellular flux analyzer, with capacitation being stimulated pharmacologically via the addition of a cAMP analog (db-cAMP) and a phosphodiesterase inhibitor (IBMX) in the absence of NaHCO_3_. In our case, sperm were preincubated for 1 hour under capacitating conditions (including NaHCO_3_) before placing them in the analyzer, and extracellular flux measurements were performed after sperm capacitation had occurred in a high proportion of cells in the population. Although pharmacological elevation of cAMP levels is sufficient to promote the protein phosphorylation events associated to capacitation in mouse sperm [30,67,77], the absence of HCO_3_^−^ might affect other capacitation-related processes that depend on the concentration of this ion [15,36,81,82], such as membrane hyperpolarization, intracellular pH increase, and hyperactivation, and could impact the activity of metabolic pathways [83–85]. In addition, the effects of sperm capacitation on sperm metabolism found by the present study are smaller in magnitude than those reported by Balbach *et al*. [68] The changes in metabolic activity following pharmacologically-stimulated capacitation ranged between 5-7 (OCR) and 4-5-fold (ECAR) in their study. In the present study, incubation under capacitating conditions resulted in a decrease of 37 % in basal respiration rate and an increment of 52 % in basal glycolysis rate. Finally, comparison between treatments in the previous study is problematic since phenomena such as non-mitochondrial oxygen consumption, and non-glycolytic acidification were not estimated by subtracting values after inhibition from basal measurements.

As an alternative to a general increase in sperm metabolism linked to capacitation [67], our results suggests that sperm capacitation promotes a change in the usage ratio of the two main metabolic pathways, from a highly oxidative metabolic phenotype in non-capacitated sperm to a predominantly glycolytic one in capacitated sperm (Fig. 7B). Since oxidative phosphorylation yields more ATP per molecule of glucose consumed than aerobic glycolysis, evidence does not support the hypothesis that sperm would be expected to reorient their metabolism to pathways that produce ATP more efficiently in response to increased ATP demands associated with capacitation [58,64]. Nonetheless, some caveats should be considered.

Recent studies have found that mouse sperm capacitation is associated with increased glucose uptake [66], and that active glycolysis is crucial for the maintenance of ATP concentration and bending capacity in the distal end of the flagellum [50]. This suggests that (a) ATP produced in the mitochondria does not diffuse sufficiently fast in order to reach the end of the flagellum, and (b) local ATP provision is a crucial physiological variable for the motility of these cells. According to our results, capacitated mouse sperm do not appear to engage in increased ATP consumption and are able to maintain ATP levels similar to those of non-capacitated cells. Considering that aerobic glycolysis occurs along most of the principal piece, a high glycolytic rate would provide a metabolic alternative to ensure sufficient ATP supply if physiological events associated with sperm capacitation (ion transport, protein phosphorylation, or hyperactivated movement) resulted in modifications of the local ATP demands in specific flagellar regions. Local ATP generation by mechanisms other than OXPHOS is relatively common in cellular structures with high energetic demands whose dimensions or structure make them unable to accommodate mitochondria [86]. In addition, it should also be borne in mind that EFA and ATP consumption analyses were performed in sperm populations with a high proportion of capacitated sperm [72] and, thus, early processes that take place during the acquisition of capacitation, and that may affect energy metabolism, have been excluded from this study.

In coincidence with previous results [67], inhibition of glycolysis resulted in a decrease of OXPHOS rates even when the culture medium was supplemented with respiratory substrates, thus indicating that the substrate used to fuel the Krebs cycle in the mitochondria of these cells is necessarily of endogenous origin. In support of this interpretation, earlier studies revealed that in a medium containing glucose, the monocarboxylate transporters expressed in the flagellar membranes of mouse sperm (MCT2) are predominantly used to export lactate instead of importing respiratory substrates [84,87,88]. In all, the alteration of the OCR/ECAR ratio promoted by capacitation might be simply regarded as an increase in the proportion of the pyruvate that, after its generation in the principal piece, is exported as lactate instead of being transported to the mitochondria to be oxidized (see Fig. 7B for a graphical conclusion). While the diversion of potential OXPHOS substrates to an excretion pathway might appear as a waste of energetic resources for the cell, the reduction of pyruvate by lactate dehydrogenase (LDH) to produce lactate involves the regeneration of the NAD+ pool (from oxidized NADH), a crucial requirement for the maintenance of high glycolytic rates [89,90].

A recent publication has shown that mouse sperm increase their mitochondrial membrane potential (MMP) during capacitation under physiological conditions [68,69], and that this event is a necessary condition to achieve hyperactivated motility [69]. These findings might appear contrary to our results, since they were first interpreted as a sign of higher OXPHOS in capacitated cells. In other words, high MMP would be associated to elevated mitochondrial ATP synthesis. This is a common misconception since H^+^ reentry through the ATP synthase during ADP phosphorylation causes a drop in MMP. Thus, higher ATP synthesis rates result in lower MMP [91]. The transition from state 3 (active ATP synthesis) to state 4 (e.g., oligomycin inhibited ATP synthesis) would in turn lead to MMP hyperpolarization [91–93]. Thus, the increased MMP exhibited by capacitated mouse sperm when compared to non-capacitated cells [68,69], would be consistent with the decrease in OXPHOS rates observed in our study.

Our study also revealed that inhibition of mitochondrial ATP production with oligomycin resulted in increased glycolytic rates. The decrease in ATP concentration produced by OXPHOS inhibition leads to the activation of the AMP protein kinase (AMPK), which, among other effects, has been shown to promote an increase in catabolic processes (i.e., glycolysis) [94]. Thus, mouse sperm are capable of shifting the burden of ATP supply to glycolysis when respiration is insufficient to meet the energetic demands of the cell. Various degrees of metabolic flexibility are observed in cell types that are frequently subjected to unpredictable environments [95,96]. In the case of mouse sperm, this would represent a significant evolutionary advantage since it would allow them to sustain an adequate ATP supply during variations in availability and concentration of metabolic substrates that are known to exist within the reproductive tracts of female mice [97,98]. On the other hand, mouse sperm seem unable to perform the reverse compensation (i.e., increasing respiratory rate when glycolytic ATP production is unavailable) since the inhibition of glycolysis by 2DOG produces a decrease in OXPHOS of a similar level than that caused by oligomycin. The inability by mouse sperm to utilize exogenous pyruvate and their dependence on glycolysis to provide endogenous respiratory substrates has been previously reported [67], and it seems to be at odds with the significant presence of pyruvate and lactate in the mouse uterine, oviductal, and follicular fluids [97,98].

In conclusion, our study has revealed that mouse sperm exhibit a shift in the usage ratio of the main metabolic pathways, from oxidative to glycolytic, as a consequence of capacitation. These changes do not seem to affect intracellular ATP concentration and ATP consumption rates. Thus, the probable function of this metabolic switch is not to increase overall ATP production, but to sustain ATP levels while ensuring the local provision of ATP to flagellar regions with specific ATP demands that arise from capacitation. The molecular mechanisms that govern and regulate these phenomena are still poorly understood, and further research is needed to address these issues.

## Acknowledgements

We are grateful to Juan Antonio Rielo for supervising animal facilities and Esperanza Navarro for animal care at the Museo Nacional de Ciencias Naturales in Madrid.

## Author Contributions

E.R.S.R., E.R., M.T., and E.S. conceived the study. E.S. and M.T. performed experiments. M.T., E.S., and E.R.S.R. analyzed data. M.T., E.S., E.R. and E.R.S.R. wrote the paper. All authors have read and agreed to the final version of the manuscript.

