## Supplementary Materials for "Capacitation promotes a shift in the energy metabolism in murine sperm"

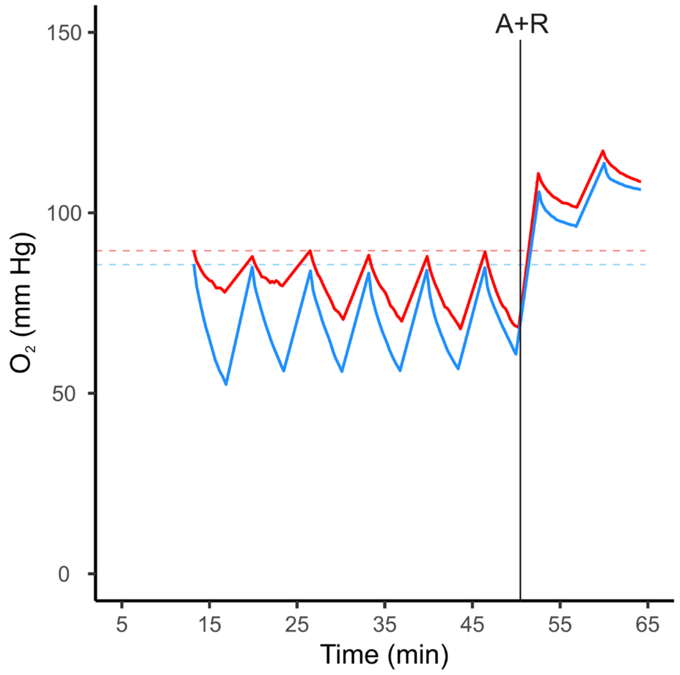


**Figure S1**. Real-time measurement of oxygen levels during extracellular flux analysis using free-swimming mouse sperm. Each oxygen curve corresponds to one well of a Seahorse XFp microplate during a representative experiment. When the sensor cartridge is lowered, forming a closed microchamber between the bottom of the plate and the cartridge, oxygen is consumed by the contained cell population (decreasing phase) and the rate of consumption is estimated by the analysis software. After measurement, the cartridge is elevated and the content of the wells are mixed, leading to the recovery of previous oxygen levels (increasing phase). Sperm were incubated in non-capacitating (blue line) or capacitated (red line) conditions for 1h previous to the experiment. Dashed horizontal lines: oxygen level at the beginning of the first measurement. Black vertical line: addition of 1 µM antimycin and 1 µM rotenone.

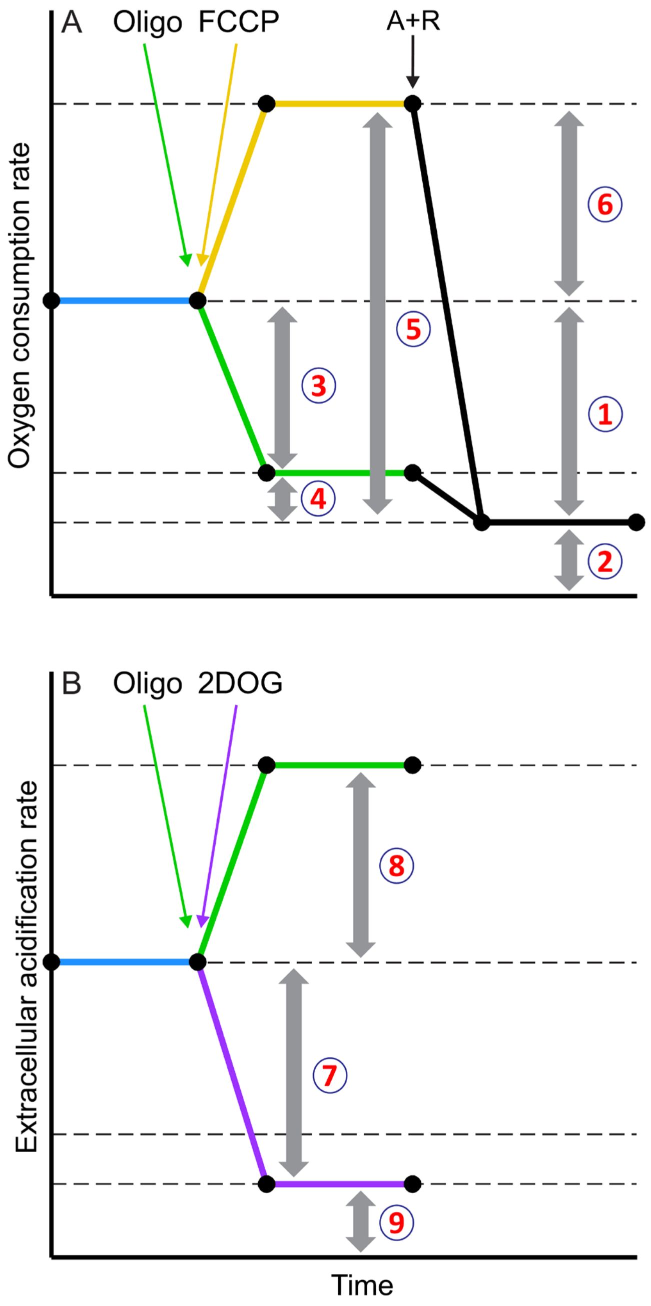


**Figure S2**. Metabolic parameters calculated to assess sperm bioenergetic phenotype. The rates of oxygen consumption (A) and extracellular acidification (B) are monitored while alternative metabolic effectors are added. Oligo: 5 µM oligomycin is added to inhibit mitochondrial ATP synthesis; FCCP: 1 µM FCCP (respiration uncoupler) is added to ensure the maximum rate of respiration is reached; A+R: simultaneous addition of 1 µM antimycin A and 1 µM rotenone to inhibit mitochondrial respiration; 2DOG: addition 50 mM 2-deoxy-glucose to inhibit glycolytic ATP and pyruvate production. The following bioenergetic parameters are calculated for the rates of oxygen consumption and extracellular acidification as indicated in the scheme: 1) Basal respiration rate; 2) Non-mitochondrial oxygen consumption; 3) Respiratory ATP production; 4) Proton leak; 5) Maximum respiration rate; 6) Spare respiratory capacity; 7) Basal glycolysis rate; 8) Glycolytic reserve; 9) Non-glycolytic extracellular acidification.

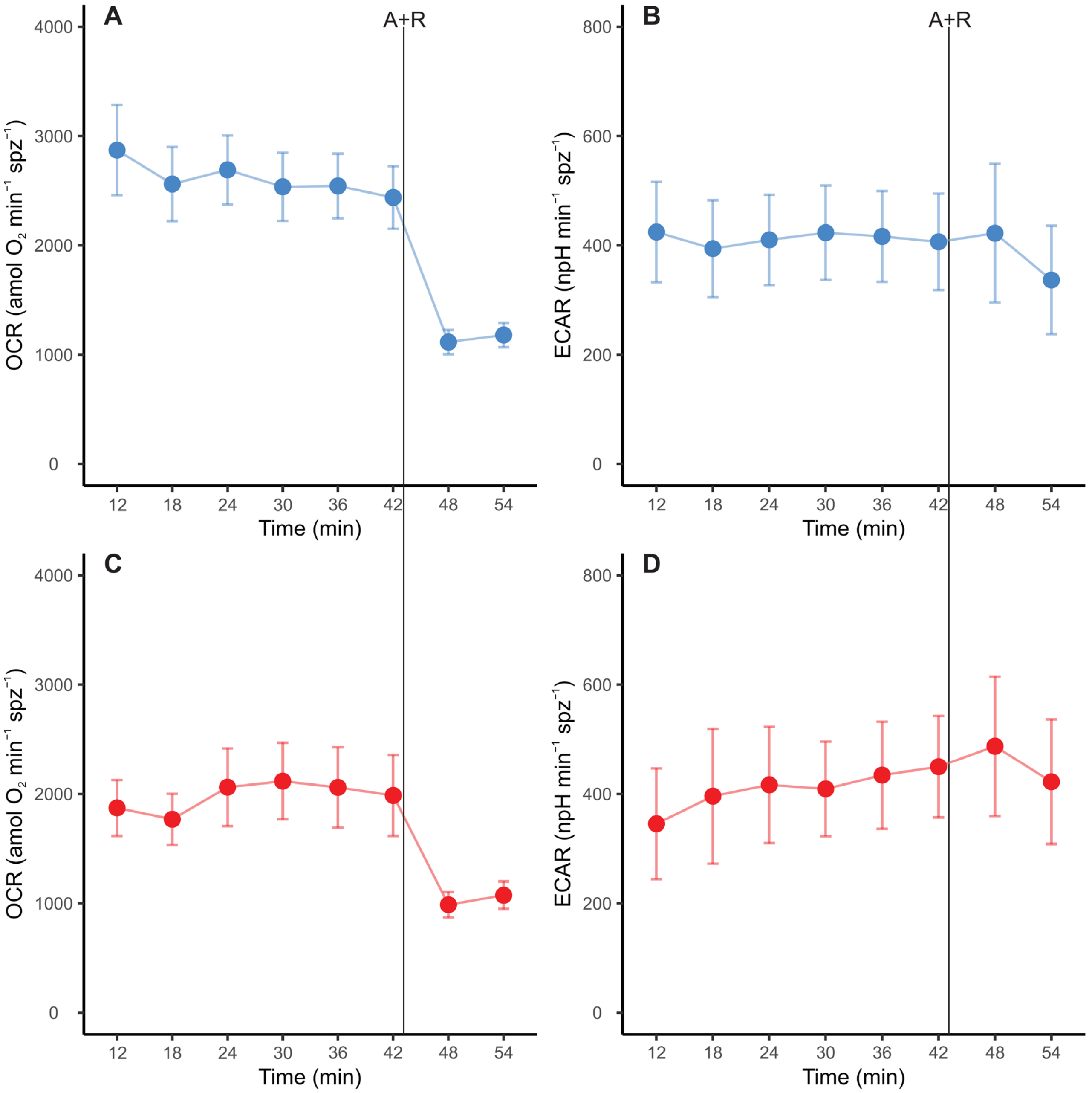


**Figure S3**. Real-time measurements of oxygen consumption rate (OCR) (A, C) and extracellular acidification rate (ECAR) (B, D) in mouse sperm. Sperm were incubated during 1 hour in non-capacitating (A, B - blue symbols) and capacitating (C, D - red symbols) conditions prior to extracellular flux analysis. Values have been normalized by sperm numbers inside each well. Symbols and whiskers correspond to means ± standard error. Time = 0 was defined as the start of the 1st measurement cycle; measurement cycles 3 to 10 are reported. The line labeled as “A+R” marks addition of 1 µM antimycin A + 1 µM rotenone.

**Table S1.** Calculation of sperm metabolic parameters based on OCR and ECAR values obtained from extracellular flux analyses.

| Parameter | Condition A | Condition B |
|---|---|---|
| Basal respiration rate[a] | OCR before any additions | OCR after the addition of antimycin A and rotenone |
| Respiratory ATP production[a] | OCR before any additions | OCR after the addition of oligomycin |
| Proton leak[a] | OCR after the addition of oligomycin | OCR after the addition of antimycin A and rotenone |
| Maximum respiration rate[b] | OCR after the addition of FCCP | OCR after the addition of antimycin A and rotenone |
| Spare respiratory capacity[b] | OCR after the addition of FCCP | OCR before any additions |
| Basal glycolysis rate[c] | ECAR before any additions | ECAR after the addition of 2DOG |
| Glycolytic reserve[a] | ECAR after the addition of oligomycin | ECAR before any additions |

Each parameter is the result of subtracting the values under condition B from the values under condition A. [a]Calculated only in experiments treated with oligomycin, antimycin and rotenone. [b]Calculated only in experiments treated with FCCP. [c]Calculated only in experiments treated with 2DOG.
